# Along- and cross-muscle fiber shear moduli in skeletal muscle

**DOI:** 10.1101/2024.07.31.605692

**Authors:** Ridhi Sahani, William E. Reyna, Thomas Royston, Eric J. Perreault, Daniel Ludvig

**Affiliations:** Biomedical Engineering, Northwestern University, Evanston IL USA; Shirley Ryan AbilityLab, Chicago IL USA; Biomedical Engineering, University of Illinois Chicago, Chicago IL USA; Physical Medicine and Rehabilitation, Northwestern University, Chicago IL USA

**Author notes:** co-first authors (Ridhi Sahani, William E. Reyna). co-senior authors (Eric J. Perreault, Daniel Ludvig). Corresponding Author: Ridhi Sahani Address: Department of Biomedical Engineering, 2145 Sheridan Road, Evanston, IL 60208.

**Keywords:** Shear Modulus, Anisotropy, Skeletal Muscle, Architecture

## Abstract

The material properties of muscle play a central role in how muscle resists joint motion, transmits forces internally, and repairs itself. While many studies have evaluated muscle’s tensile material properties, few have investigated muscle’s shear properties. The objective of this study was to quantify the shear moduli of skeletal muscle both along (along-muscle fiber) and perpendicular (cross-muscle fiber) to the direction of muscle fibers. We collected data from the extensor digitorum longus, tibialis anterior, and soleus muscles harvested from both hindlimbs of 12 rats. These muscles were chosen to further evaluate the consistency of shear moduli across muscles with different architectures. We applied strains and measured stress in three configurations: parallel, perpendicular, and across the muscle fibers to characterize the along- and cross-muscle fiber tensile and shear material parameters. We found no significant difference between the shear modulus measured parallel to the fibers (along-muscle fiber) and the shear modulus in the plane perpendicular to the fibers (cross-muscle fiber). Although the shear moduli were not significantly different, there was a greater difference with increasing strain, suggesting that there is greater anisotropy at larger strains. We also found no significant difference in moduli between the muscles with differing muscle architecture. These results characterize the shear behavior of skeletal muscle and are relevant to understanding the role of shear in force transmission and injury.

## Introduction

The mechanical properties of skeletal muscle are essential for the generation of forces required for human movement. Changes in muscle properties, such as increased stiffness, contribute to dysfunction and impairment across numerous neuromuscular disorders and during injury. To improve our understanding of and treatment for such conditions it is important to develop methods to characterize muscle properties, but this remains challenging due to the complex three-dimensional geometries, loading states, and force transmission pathways in skeletal muscles. Active forces, generated by contractile elements, and passive forces, generated by passive structures such as the extracellular matrix (ECM), are transmitted longitudinally along the length of muscle fibers, and laterally between neighboring muscle fibers. Longitudinal force transmission occurs directly through the myotendinous junction and is often considered to be the main pathway. However, experimental studies in which the longitudinal pathway is physically disrupted show that the majority of contractile force is transmitted laterally.^1,2^ Three-dimensional models reveal the mechanism of lateral force transmission to be shearing between muscle fibers and surrounding ECM, which is essential for force transmission between intrafascicularly terminating fibers.^3^ Both shear and tensile deformations occur within skeletal muscles, and shearing is a common mode of muscle injury, causing rupture of myofibers, surrounding connective tissues, and blood vessels^4^. The shear modulus of skeletal muscle, or its resistance to shear stress, is an important material property due to the role of shear loading on force transmission and injury.

The shear properties of skeletal muscles are not well understood. While the tensile stiffness (young’s modulus) of muscle has been extensively characterized,^5–8^ studies in which shear properties are characterized are limited.^7,9,10^ Material properties along the direction of muscle fibers are typically measured, but skeletal muscle is considered to be transversely isotropic, ^11^ where material properties parallel to the muscle fibers (along-muscle fiber) differ from those in the plane of isotropy perpendicular to the fibers (cross-muscle fiber). Morrow et al measured shear in the along-muscle fiber direction in rabbit extensor digitorum longus muscle^7^ as well as the Young’s modulus in along-and cross-muscle fiber directions. These three measurements were sufficient for characterizing the material properties of a transversely isotropic material^12^ but lacked independent verification on the cross-muscle fiber direction and were also restricted to a single muscle architecture. Moriera et al applied shear loading in bovine sternomandibularis muscle with fibers oriented at 0°, 45°, or 90° relative to the applied load and with varying preloads.^10^ They measured greater stresses during longitudinal shear in the direction of muscle fibers compared with shear in the plane perpendicular to muscle fibers, highlighting the importance of measuring shear properties in multiple planes.

Previous experiments have been limited to a single muscle group and it is important to evaluate the consistency of shear modulus measures across muscles with differing architectures. Differences in architectural features such as physiological cross-sectional area (PCSA), pennation angle (α), and fiber lengths (l_f_) are seen across skeletal muscles.^13^ These architectural features are related to the functional demands of specific muscles and influence the distribution of loads^14^ and nonuniformity of strains^11^ within muscle. For example, the pennation angle, a measure of muscle fiber orientation relative the muscle line of action, can result in varying levels of force generation^15^ and transmission.^16^ The shear modulus is implicated on the resistance to loading, but it is unknown if and how differences in shear properties relate to architectural differences between muscles. Therefore, it is important to consider if and how along-and cross-muscle fiber shear moduli vary between skeletal muscles.

The objective of this study was to quantify both the along-muscle fiber and cross-muscle fiber shear moduli of skeletal muscle tissue. Our primary hypothesis was that the shear modulus would differ within the different planes of shear relative to the orientation of the muscle fibers. We also hypothesized that the shear modulus would not differ across muscle types, based on the assumption that material properties are consistent between muscles and differences arise due to their unique geometries and architectural features. We tested these hypotheses by measuring the shear modulus in the extensor digitorum longus (EDL), tibialis anterior (TA), and soleus (SOL) muscles from rat hindlimbs. Measures were made at three orientations: parallel, perpendicular, and across the muscle fibers and used to characterize the along- and cross-muscle fiber tensile and shear material parameters. Our results provide values for both the along-muscle fiber and cross-muscle fiber shear moduli of skeletal muscle, providing insight to this important mechanical property and values that can be used to simulate muscle responses through continuum modeling. This is an important step towards evaluating shearing forces in muscle and further understanding the role of muscle material properties in force transmission and muscle injury.

## Materials and Methods

### Tissue Collection and Preparation

All data were collected from healthy Sprague-Dawley rats approximately 15-month-old (n=12 rats) obtained through the Northwestern University tissue sharing program. The EDL, TA, and SOL muscles were harvested from both hindlimbs immediately following termination and placed in chilled phosphate-buffered saline to stem the effects of rigor.^17^ Muscles had their aponeuroses dissected away using a surgical scalpel to ensure measurements were made only on muscle tissue, similar in procedure to Morrow.^7^ Before mechanical testing, muscles were sectioned under a microscope into rectangular cubes (∼9 x 9 x 4 mm) aligned with the muscle fibers (Fig 2A) . The length, width, and thickness of each specimen (rectangular cube) was measured while the tissue was physically unconstrained. Multiple specimens were obtained from some muscles, resulting in a total of 106 specimens. Shear measurements were made with the cubes oriented in one of three directions: parallel, perpendicular, or across muscle fibers (Fig. 1). Specimens (24 total) were eliminated if they showed separation from our testing apparatus during testing or if artifacts, such as transient changes in force, were observed in the recordings.

**Figure 1.**
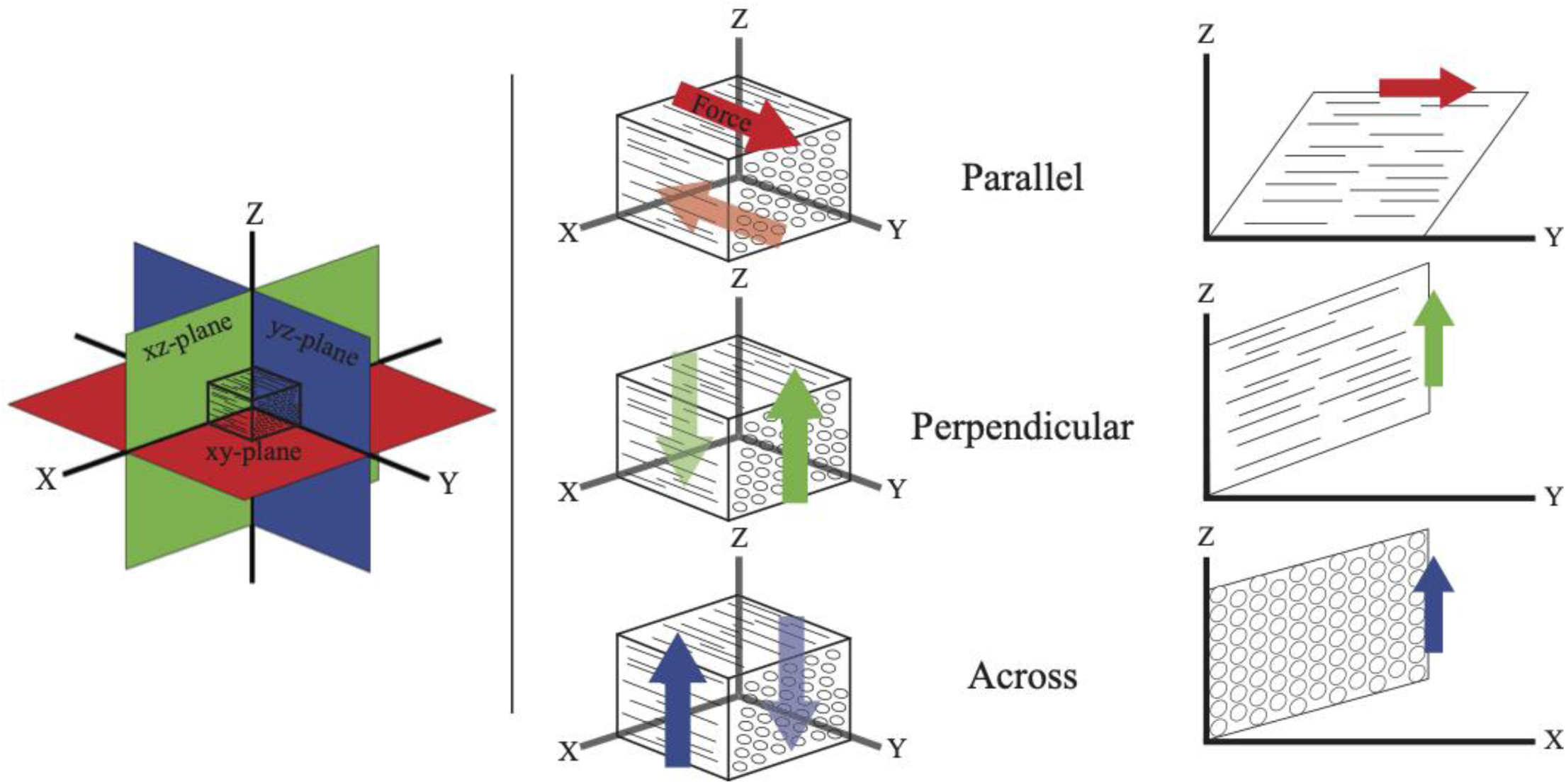
Three-dimensional representation of shearing muscle in the parallel (xy), perpendicular (xz), and across (yz) planes. Solid arrows indicate direction of force and displacement, while transparent arrows indicate stationary resistant force.

**Figure 2.**
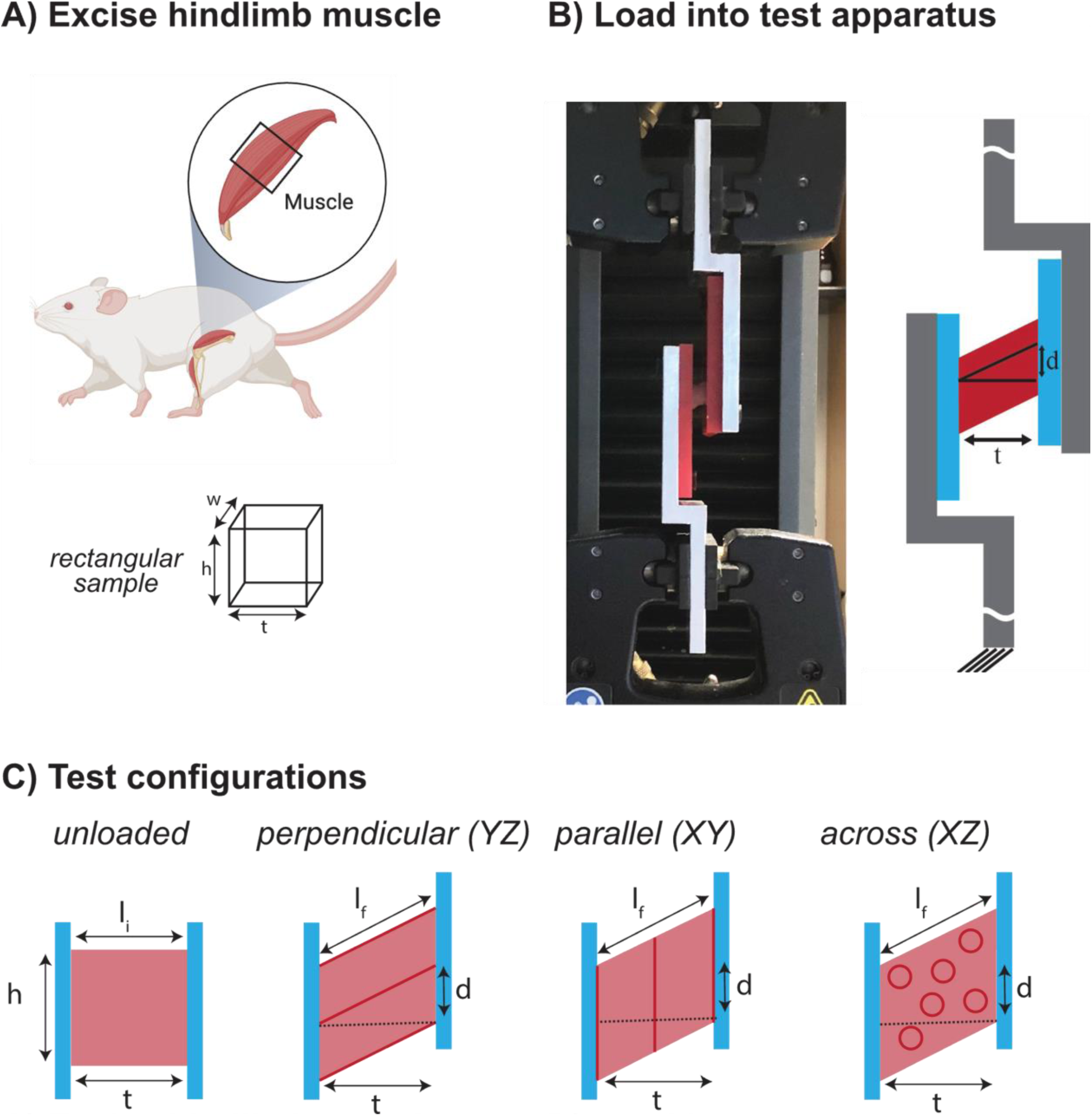
Overview of experimental methods. A) Rectangular sample excised from hindlimb muscles, B) shear testing apparatus photograph (left) and schematic (right). Exaggerated representations of the thickness (t), displacement (d), and force (F) are show in the schematic. C) Test configurations for shear loading in the across, parallel, and perpendicular planes.

### Mechanical Testing

Muscle specimens were tested within 12 hours of harvesting. Displacements were applied using an Instron mechanical tester (Instron 5942; Instron Corp., Canton, MA). An offset aluminum bracket with acrylic inserts was mounted into the uniaxial tensile tester, similar in design to Morrow^7^ (Fig. 2B). Aluminum brackets were milled in unison to ensure that surfaces were parallel to one another. Acrylic inserts were used to promote adhesion of the tissue and to easily exchange the testing surface between specimens. The harvested muscle specimens were fixed to the acrylic plates using cyanoacrylate glue. The plates were then screwed onto the aluminum brackets to form a rigid clamp. Specimens were positioned in the center of the testing apparatus to ensure that forces were uniformly applied to the tissue faces and that measurements were in line with the load cell. The use of cyanoacrylate glue bonded tissue to the fixture beyond muscle failure, as seen by rupture within samples as opposed to separation from our mounting plates. The specimen height was remeasured in the clamp to replicate the stress-free state on the bench. Data were collected at a strain rate of 5% per minute until failure, as indicated by a sudden drop in force and tearing of the muscle.

### Calculation of material properties

For all calculations, second Piola-Kirchhoff stress was used, referencing the undeformed state of a material with a constant cross-section. Our cubic specimens, to the best of our abilities, had a constant cross-section and our adhered surface encompassed the entire face of the specimen. Green shear strain was used because of the small deformations in length. The measured stress was calculated as the applied force (*F*) divided by the initial cross-sectional area (Fig. 2B).

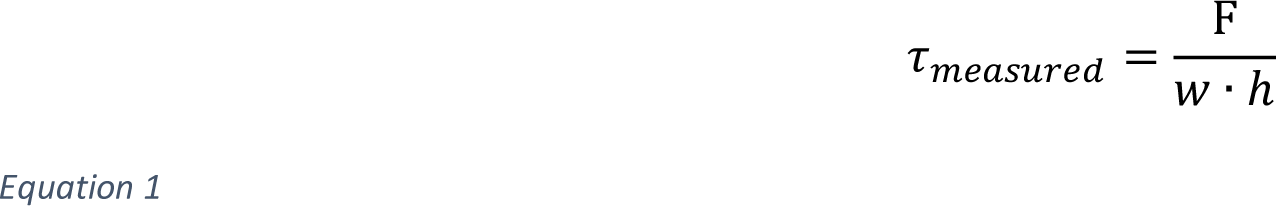

The applied shear strain (γ) was calculated as the inverse tangent of the applied displacement (*d*) divided by the thickness (*t*) of the specimen.

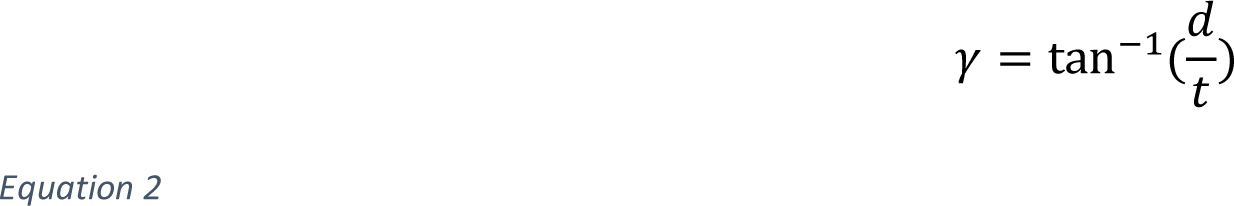

For feasibility of testing, one plate was fixed and a vertical displacement was applied to the other plate in our experimental setup. Therefore, a normal strain (*ε*) was also applied to the specimen and can be calculated based on the change in the length of the specimen, ε = (ℓ_*f*_ − *t*)/*t*, or expressed in terms of γ:

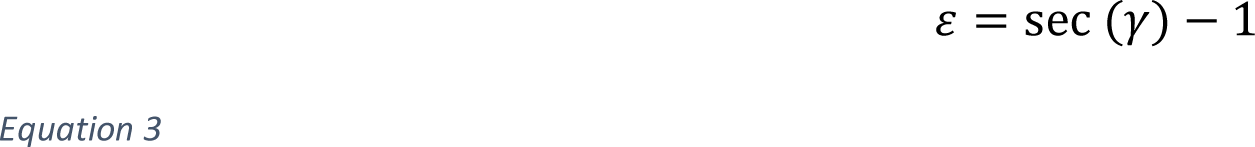

As a result the stress measured by the Instron included both a normal and shear component. The normal stress (*σ*) can be calculated based on the young’s modulus (*E*) and component of normal strain parallel to the applied displacement direction,

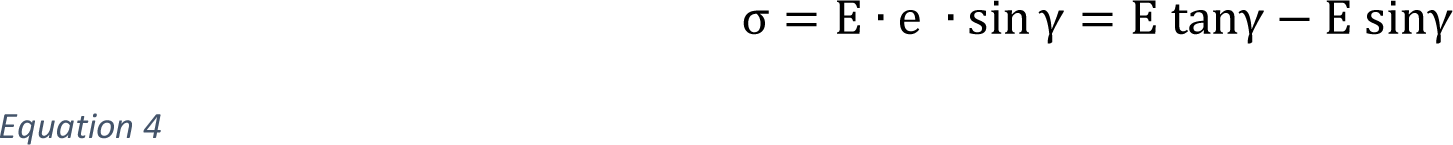

while the shear stress (*τ*) can be calculated based on the shear modulus (*µ*) and the applied shear strain.

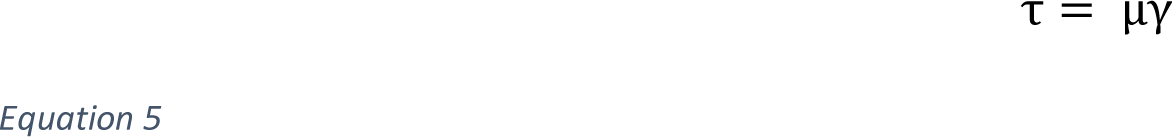

Therefore, the total stress measured by the Instron is

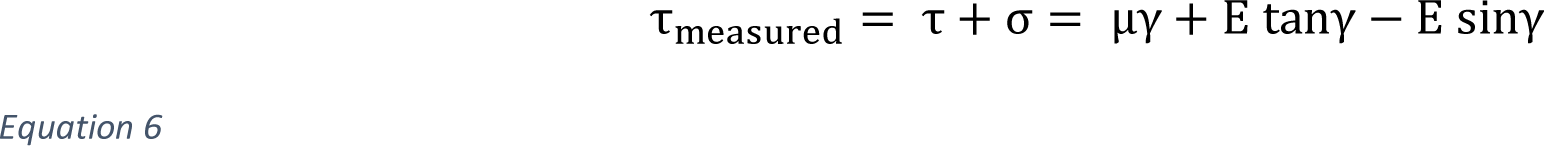

and the instantaneous slope (k) can be determined by taking the first derivative of τ_measured_ with respect to γ.

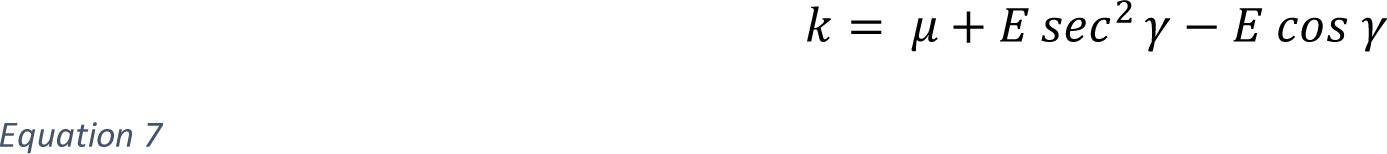

The orientation of the muscle fibers relative to the applied displacement varied between test conditions, with unique combinations of shear and normal stresses and strains in the along- and cross-muscle fiber directions. Therefore, the instantaneous slopes of the measured stress-strain relationships for the parallel (*k_XY_*), perpendicular (*k_YZ_*), and across (*k_XZ_*) conditions can be expressed as:

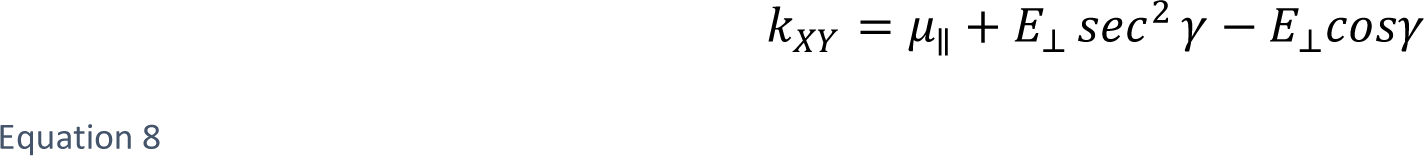

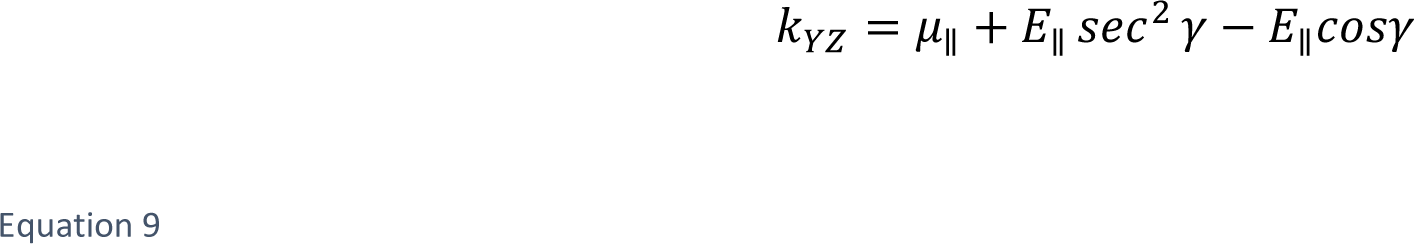

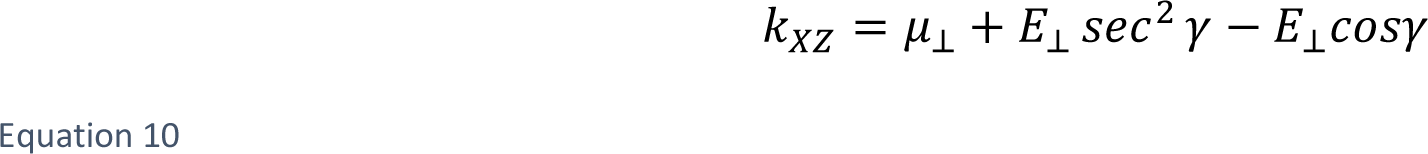

and depend on four material parameters: along-muscle fiber shear modulus (μ_∥_), cross-muscle fiber shear modulus (μ_⊥_), along-muscle fiber young’s modulus (*E*_∥_), and cross-muscle fiber young’s modulus (*E*_⊥_).

For a linear elastic nearly incompressible, transversely isotropic material, only three of the four parameters must be defined to fully describe the material. Therefore, we can use the following relationship :

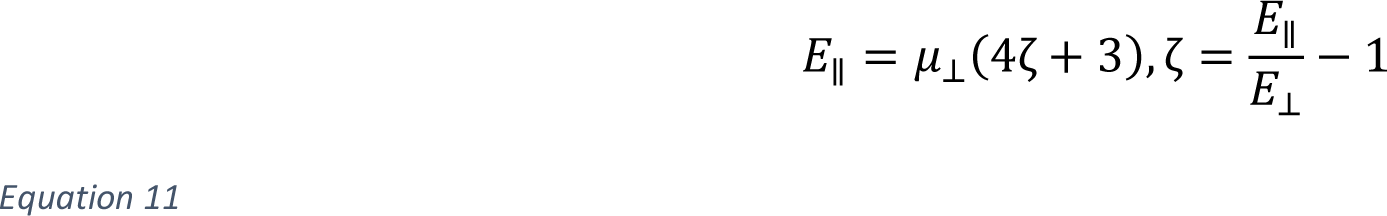

along with equations 8-10 to solve a system of equation with four unknown parameters.

The raw load-extension curves from each test were down sampled, adjusted for machine error, clipped at failure, and filtered. The measured stress and applied strains were calculated using Eqns 1-2. The instantaneous slope of the measured stress-applied strain relationship (*k*) was determined at strain intervals of 0.02 between the range of strains of 0.1 to 0.4, which falls within the predicted physiological range for along-muscle fiber and cross-muscle fiber shear strains (Fig. 3).^11^ A linear mixed-effects model was used to estimate the instantaneous slopes across the three test conditions (parallel, perpendicular, across) and the three different muscles (EDL, SOL, TA). We fitted the model with instantaneous slope as the dependent variable, strain, muscle type and orientation as fixed factors, and specimen as a random factor. The values for each instantaneous slope were estimated at each strain value and used to solve for µ_II_, µ_T_, E_II_, and E_T_ using equations 8-11 (MATLAB, Symbolic Math Toolbox). The standard error for each of the 4 moduli were estimated using the standard errors returned by linear mixed-effects model.

**Figure 3.**
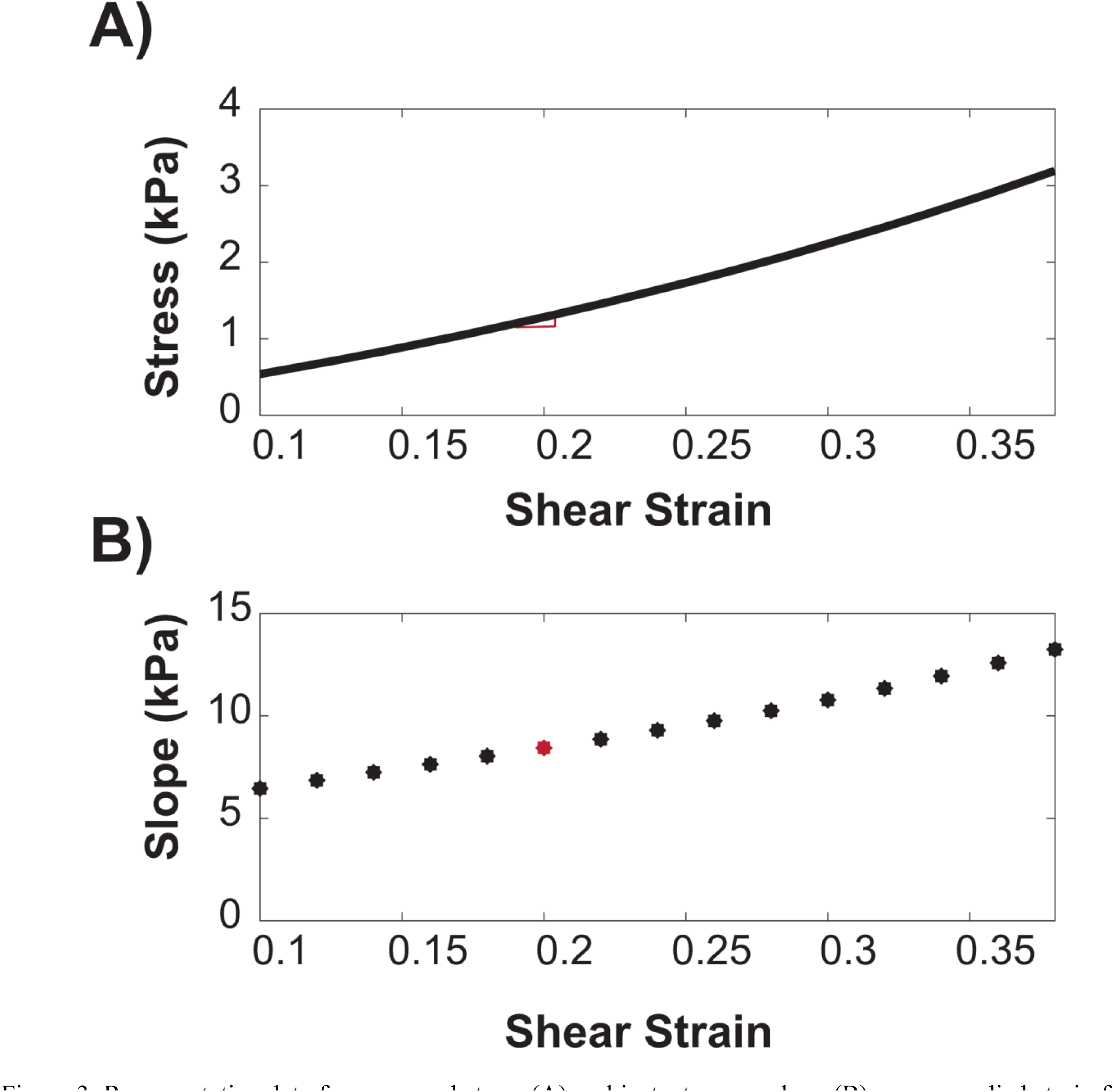
Representative data for measured stress (A) and instantaneous slope (B) versus applied strain from one soleus muscle specimen in the parallel configuration. Circles indicate the strain values selected within the physiological strain limit at which the instantaneous slope was calculated of the shear stress-strain relationship, whose calculation for a single point is denoted by the red triangle.

### Statistical Analysis

Our primary hypothesis was that the shear and young’s moduli would differ between the along-muscle fiber and cross-muscle fiber directions. We tested this hypothesis using a Wald test with Satterthwaite approximation^19^ at each strain level for each of the moduli. Our secondary hypothesis was that the material properties would be consistent across the different muscle types. We tested this hypothesis using a Wald test with Satterthwaite approximation^19^ at each strain level for each of the material properties. All analyses were carried out in Matlab (Mathworks, Natick, MA). Results are presented as mean ± standard error, unless otherwise specified.

## Results

For all muscles grouped together, the measured stress-strain slope was greater in the perpendicular compared with the parallel and across configurations (p<0.042) (Fig. 4). The parallel and across configurations were not significantly different from each other (p>0.83). In the parallel configuration, the measured stress-strain slope ranged from 2.87±0.51kPa to 5.43±0.52kPa. In the perpendicular configuration, the measured stress-strain slope ranged from 4.33±0.65kPa to 10.03±0.65kPa. In the across configuration, the measured stress-strain slope ranged from 2.12±0.56kPa to 4.15±0.56kPa.

**Figure 4:**
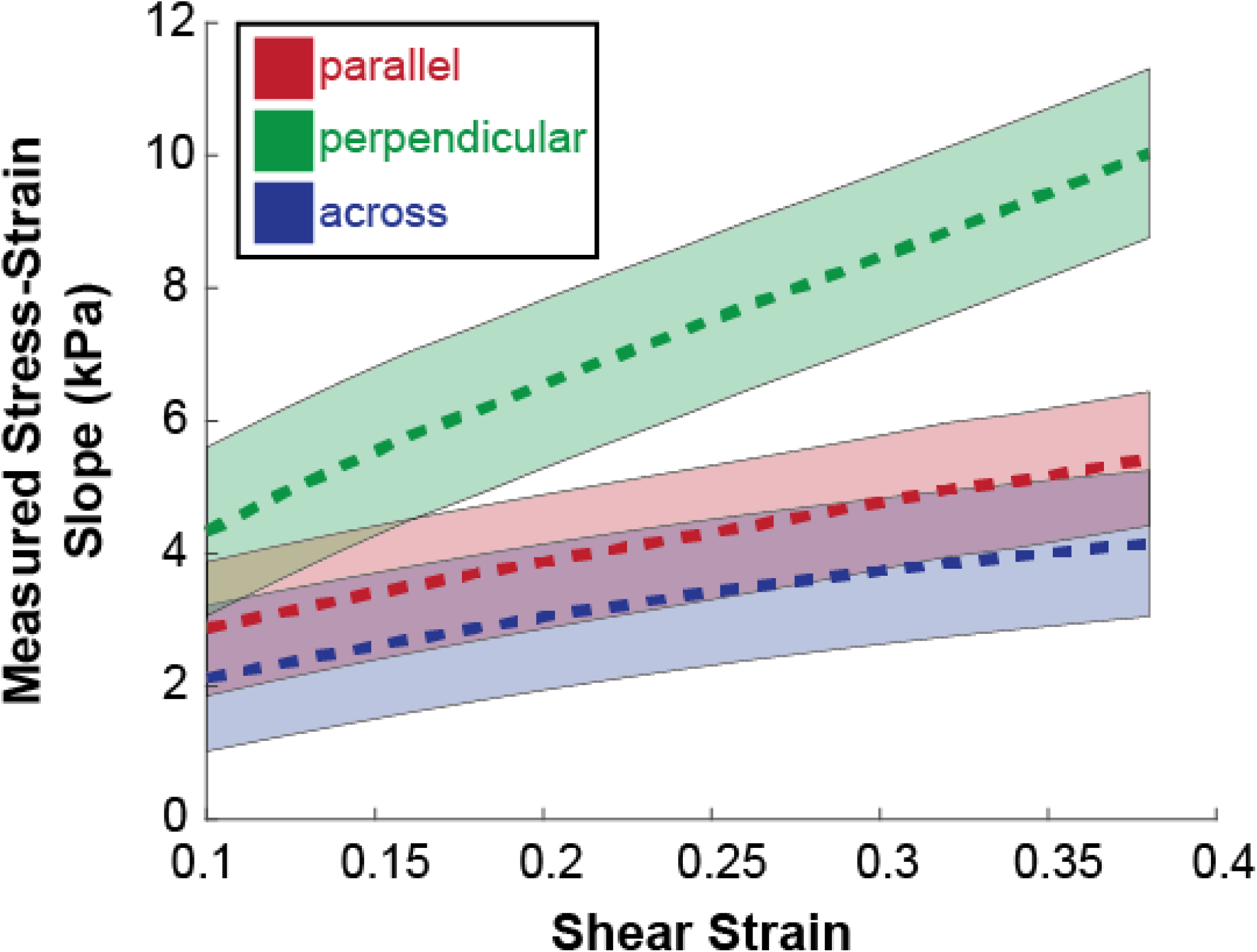
Instantaneous slopes from relationship between measured stress and applied shear strains in the parallel (red), perpendicular (green), and across (blue) conditions. Dashed lines represent mean and shaded area represents 95% confidence interval.

After accounting for the normal and shear stress components in each configuration, we calculated the along- and cross-muscle fiber shear and young’s moduli for all muscles grouped together (Fig. 5). The cross-muscle fiber shear moduli ranged from 1.69±0.50kPa to 1.30±0.26kPa over the range of applied shear strains. The along-muscle fiber shear moduli ranged from 2.44±0.58kPa to 2.58±0.66kPa over the range of applied shear strains. Both shear moduli were fairly constant over the range of shear strains. Due to the large variability in measurements between samples, there was no significant difference (p>0.05 for all strain values) between the along- and cross-shear moduli despite the along-muscle fiber shear moduli being aproximately 75% larger than the cross-fiber moduli. At low strain values, the standard errors for the young’s moduli were quite large as evident in Fig 5B. The standard errors decreased with increasing strain and at strain of 0.38 the along-muscle fiber young’s modulus of 32.26± 4.21kPa was significantly larger the cross-muscle fiber young’s modulus of 12.34±1.43kPa (p =6.65e-07).

**Figure 5:**
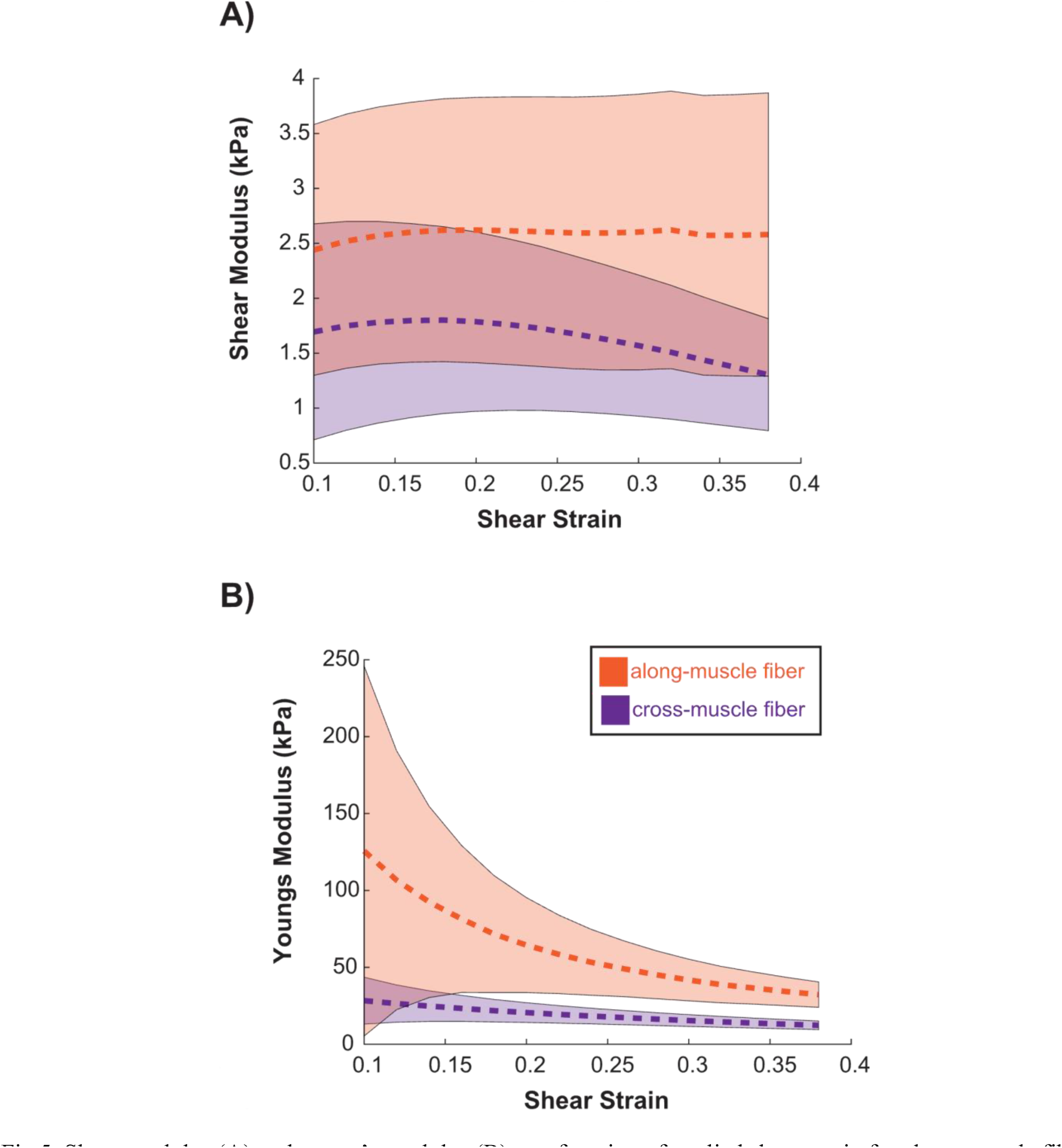
Shear modulus (A) and young’s modulus (B) as a function of applied shear strain for along-muscle fiber (orange) and cross-muscle fiber (purple) directions. Dashed lines represent mean and shaded area represents 95% confidence interval.

We did not see any significant differences in material properties between the three different muscle groups. There was a significant difference between muscle groups seen in the measured stress-strain slope in the perpendicular configuration, between the EDL and TA (p<0.015). However, this difference in slope did not result in a significant difference in any of the estimated material properties (p> 0.553) (Fig. 6). While we did not see any statistical differences between the muscles, due to the large variability in the samples, we could not conclusively demonstrate that the material properties of all 3 muscles are equivalent to any reasonable threshold.

**Figure 6:**
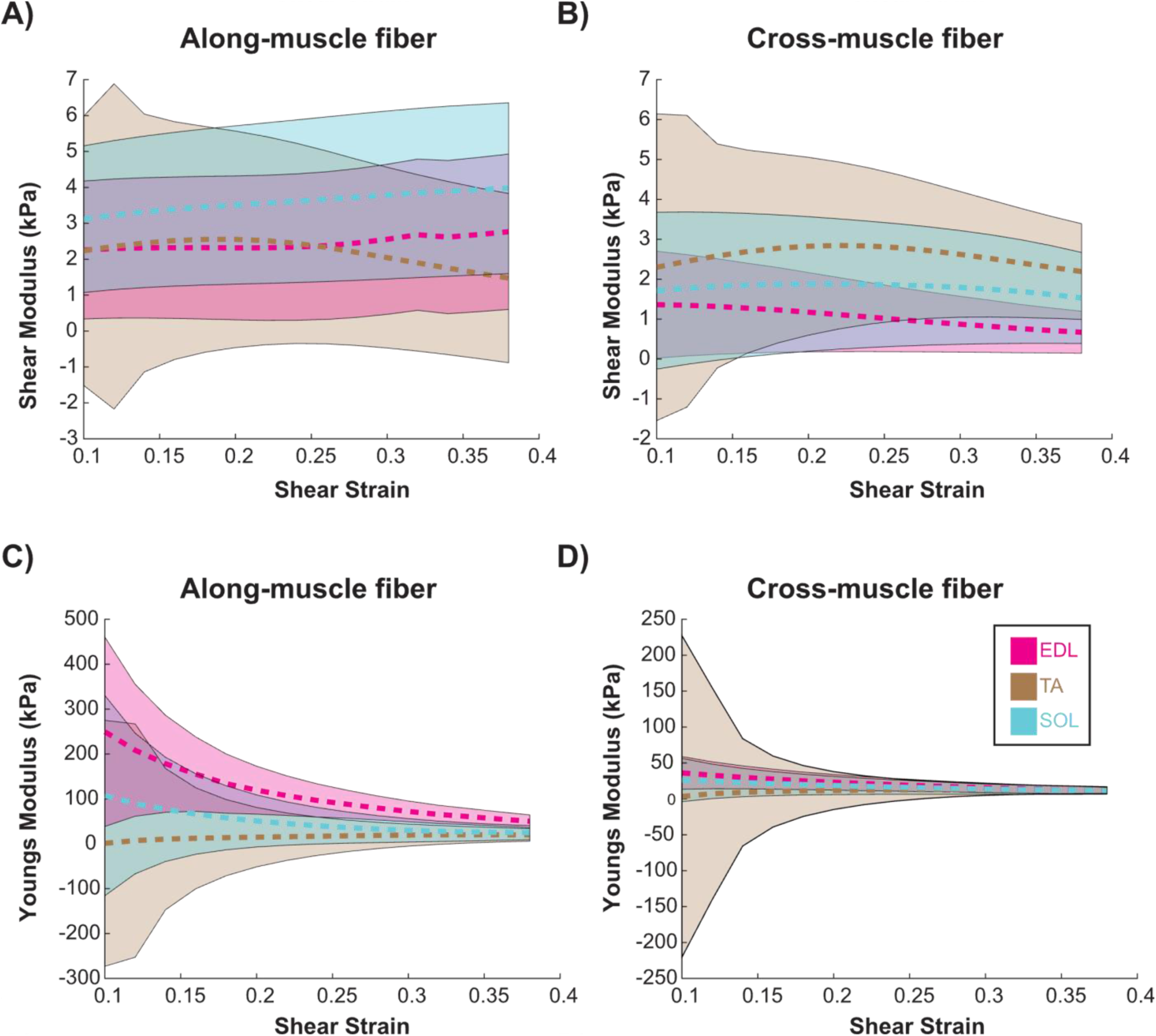
Calculated shear (A-B) and young’s (C-D) moduli for EDL (pink), TA (tan), and SOL (blue) musles. Dashed lines represent mean and shaded area represents 95% confidence interval.

## Discussion

The objective of this study was to quantify the passive shear moduli of skeletal muscle in the planes parallel and perpendicular to the direction of muscle fibers. We performed direct mechanical testing on tissue specimens from three muscles of differing architecture in the rat hindlimb. We found no significant difference between the shear modulus measured parallel to the fibers (along-muscle fiber) and the shear modulus in the plane perpendicular to the fibers (cross-muscle fiber) (Fig. 5). Although the shear moduli were not significantly different, there was a greater difference with increasing strain, suggesting that there is greater anisotropy at larger strains. We also found no significant difference in moduli between the muscles with differing muscle architecture (Fig. 6), suggesting that differences between muscles arise due to their unique geometries and architectural features rather than differences in material properties. These results characterize the shear behavior of skeletal muscle and are relevant to understanding the role of shear in force transmission and injury.

Our measurements of both along- and cross-muscle fiber shear modulus fell within the range of measurements of along-muscle fiber shear modulus from Morrow et al (3.87 ± 1.38 kPa).^7^ Our measurements of cross-muscle fiber young’s modulus were within the range of their measurements (22.4± 6.00 kPa) while our measurements of along-muscle fiber young’s modulus were below their range of measurements (447± 39.9 kPa). Their study strained samples until failure (∼40% tensile strain), while our experimental setup applied low tensile strain to the samples (<9%). As muscle exhibits a nonlinear stress-strain relationship our measures are likely in the nonlinear portion, explaining this discrepancy in measurements. Their study did not directly measure the cross-muscle fiber shear modulus, but based on their measurements and using Eqn 11, we can estimate a cross-muscle fiber shear modulus of 5.67 ± 1.68 kPa, with the standard error calculated from propagation of uncertainties assuming variables are uncorrelated. While this is greater than our measurements of cross-muscle fiber shear modulus, there is a large variability when estimating shear moduli from three other measures.

Although the shear moduli were not significantly different, the along-muscle fiber shear modulus was greater than the cross-muscle fiber shear modulus and approached a significant difference at greater strains. Similarly, Moriera et al. measured greater shear stress in the plane parallel compared with perpendicular to muscle fibers only at values above 0.5 shear strain,^10^ which was outside the range of applied strains in our experiment. We selected the range of applied shear strains for our experiment based on predictions of *in vivo* strains from computational models. Therefore, there may be minimal differences in along- and cross-muscle fiber shear moduli during typical daily movement. However, these differences may be relevant at greater levels of strain when injuries are sustained. This anisotropy is likely due to the complex organizational structure of muscle. Muscle is composed of two macroscopic structures with distinct orientations, muscle fibers and the surrounding extracellular matrix.^20–22^ The extracellular matrix is a key contributor to passive muscle stiffness^8,23^ and alterations in extracellular matrix organization are related to changes in tensile properties during disease,^24,25^ but their influence on shear properties is unknown. While we did not detect differences in shear properties across muscles, there may be disease dependent differences due to changes in extracellular matrix organization. Our methods can be applied to compare shear moduli between healthy and diseased muscles in future studies.

Our measurements can also be used to compare with ultrasound elastography estimates of shear modulus in muscle. Ultrasound elastography is a promising tool to measure in vivo muscle stiffness noninvasively based on the propagation of shear waves, ie. shear wave velocity. Shear moduli relates to shear wave velocity in isotropic materials, but this relationship is more complex in muscle due to its heterogeneity. Previous studies show that shear wave velocity is dependent on muscle activation and varies across muscles.^26,27^ Therefore, it is important to compare direct measurements of shear moduli with estimates from shear wave velocity. Previous studies in cats,^27^ pigs^28^ and humans^29–31^ exhibited shear wave speeds ranging from 2 – 10 m/s, translating to shear moduli in the range of 4 kPa to 225 kPa. Our measurements fell below this range but may be due to differences in animal models, as our data was collected in rat muscles. Future work should directly compare shear wave velocities and shear modulus in the same muscle samples.

### Limitations

While our experiment was similar in design to previous studies used to measure shear properties,^7,10^ a normal strain was also applied to the sample during shearing. Our calculations of shear moduli accounted for the normal stress component but required measurements from three different test configurations and a defined relationship between transversely isotropic material properties. Therefore, our calculations are sensitive to more sources of error and rely on the assumption of transverse isotropy, which is challenged as the muscle-tissue scale.^32^ For more direct measurements, future experiments should be designed to apply shear strains only. Our study did not characterize viscoelastic material properties, which may be important for understanding the material properties of muscle relevant to more dynamic conditions. However, we selected a slow rate of extension to eliminate the viscoelastic effects of muscle from our measures. Studies at higher rates, suitable for estimating viscoelastic properties, could be used to supplement the measures reported here. All samples were cut into cubes by hand, which may have introduced slight variations in shape or orientation relative to the muscle fibers. This was mitigated to the best of our ability by taking dimensional measures before testing and by using a large sample size. Still, this manual approach may have contributed to the large variability in our measures. The attachment of tissue samples to the clamps may have also introduced artifacts in our measurements. We used applied strain for our calculations and tracking the deformation of the tissue through methods such as digital image correlation would provide a better metric of tissue strains.

### Conclusions

This study presents measures of both the along-muscle fiber and cross-muscle fiber shear moduli in skeletal muscle tissue. The along-muscle fiber young’s modulus was greater than the cross-muscle fiber young’s modulus over the range of strain applied in our experiment. Although the along-muscle fiber shear modulus was not significantly different than the cross-muscle fiber shear modulus, greater differences were seen at larger strains. Despite differences between muscles, we did not find the shear modulus of the rat muscles we tested to be different. The quantitative measures reported here can be used to describe the mechanical properties of muscle in shear, which adds to the growing knowledge of muscle material properties as are needed to develop computational models of how muscle responds to the complex stresses and strains essential to its use in the control of movement and posture.

## Acknowledgments

This work was supported by National Institute of Health grants R01AR071162 and T32HD07418.

## References

1. Huijing, P., Baan, G. C. & Rebel, G. T. Non-myotendinous force transmission in rat extensor digitorum longus muscle. J. Exp. Biol. 201, 683–691 (1998).

2. Street, S. F. Lateral transmission of tension in frog myofibers: A myofibrillar network and transverse cytoskeletal connections are possible transmitters. J. Cell. Physiol. 114, 346–364 (1983).

3. Sharafi, B. & Blemker, S. S. A mathematical model of force transmission from intrafascicularly terminating muscle fibers. J. Biomech. 44, 2031–2039 (2011).

4. Järvinen, T. A., Järvinen, M. & Kalimo, H. Regeneration of injured skeletal muscle after the injury. Muscles Ligaments Tendons J. 3, 337–345 (2014).

5. Bosboom, E. M. et al. Passive transverse mechanical properties of skeletal muscle under in vivo compression. J. Biomech. 34, 1365–1368 (2001).

6. Lieber, R. L. & Fridén, J. Muscle contracture and passive mechanics in cerebral palsy. J. Appl. Physiol. 126, 1492–1501 (2019).

7. Morrow, D. A., Haut Donahue, T. L., Odegard, G. M. & Kaufman, K. R. Transversely isotropic tensile material properties of skeletal muscle tissue. J. Mech. Behav. Biomed. Mater. 3, 124–129 (2010).

8. Reyna, W. E., Pichika, R., Ludvig, D. & Perreault, E. J. Efficiency of skeletal muscle decellularization methods and their effects on the extracellular matrix. J. Biomech. 110, 109961 (2020).

9. Hashemi, S. S., Asgari, M. & Rasoulian, A. An experimental study of nonlinear rate-dependent behaviour of skeletal muscle to obtain passive mechanical properties. Proc. Inst. Mech. Eng. [H] 234, 590–602 (2020).

10. Moreira, C. S. & Nunes, L. C. S. Mechanical behavior of skeletal muscles under simple shear at large strain. J. Braz. Soc. Mech. Sci. Eng. 44, 511 (2022).

11. Blemker, S. S., Pinsky, P. M. & Delp, S. L. A 3D model of muscle reveals the causes of nonuniform strains in the biceps brachii. J. Biomech. 38, 657–665 (2005).

12. Crutison, J., Sun, M. & Royston, T. J. The combined importance of finite dimensions, anisotropy, and pre-stress in acoustoelastography. J. Acoust. Soc. Am. 151, 2403–2413 (2022).

13. Gans, C. Fiber architecture and muscle function. Exerc. Sport Sci. Rev. 10, 160–207 (1982).

14. Burkholder, T. J., Fingado, B., Baron, S. & Lieber, R. L. Relationship between muscle fiber types and sizes and muscle architectural properties in the mouse hindlimb. J. Morphol. 221, 177–190 (1994).

15. Lieber, R. L. & Fridén, J. Functional and clinical significance of skeletal muscle architecture. Muscle Nerve 23, 1647–1666 (2000).

16. Huijing, P. A. Muscle as a collagen fiber reinforced composite: a review of force transmission in muscle and whole limb. J. Biomech. 32, 329–345 (1999).

17. Tuttle, L. J., Alperin, M. & Lieber, R. L. Post-Mortem Timing of Skeletal Muscle Biochemical and Mechanical Degradation. J. Biomech. 47, 1506–1509 (2014).

18. Schmidt, J. L. et al. Magnetic resonance elastography of slow and fast shear waves illuminates differences in shear and tensile moduli in anisotropic tissue. J. Biomech. 49, 1042–1049 (2016).

19. Luke, S. G. Evaluating significance in linear mixed-effects models in R. Behav. Res. Methods 49, 1494–1502 (2017).

20. Purslow, P. P. & Trotter, J. A. The morphology and mechanical properties of endomysium in series-fibred muscles: variations with muscle length. J. Muscle Res. Cell Motil. 15, 299–308 (1994).

21. Trotter, J. A. & Purslow, P. P. Functional morphology of the endomysium in series fibered muscles. J. Morphol. 212, 109–122 (1992).

22. Eng, C. M. et al. Scaling of muscle architecture and fiber types in the rat hindlimb. J. Exp. Biol. 211, 2336–2345 (2008).

23. Gillies, A. R. & Lieber, R. L. Structure and function of the skeletal muscle extracellular matrix. Muscle Nerve 44, 318–331 (2011).

24. Sahani, R., Hixson, K. & Blemker, S. S. It’s more than the amount that counts: implications of collagen organization on passive muscle tissue properties revealed with micromechanical models and experiments. J. R. Soc. Interface 21, 20230478 (2024).

25. Sahani, R., Wallace, C. H., Jones, B. K. & Blemker, S. S. Diaphragm muscle fibrosis involves changes in collagen organization with mechanical implications in Duchenne muscular dystrophy. J. Appl. Physiol. 132, 653–672 (2022).

26. Bernabei, M., Lee, S. S. M., Perreault, E. J. & Sandercock, T. G. Axial stress determines the velocity of shear wave propagation in passive but not active muscles in vivo. J. Appl. Physiol. 134, 941–950 (2023).

27. Bernabei, M., Lee, S. S. M., Perreault, E. J. & Sandercock, T. G. Shear wave velocity is sensitive to changes in muscle stiffness that occur independently from changes in force. J. Appl. Physiol. 128, 8–16 (2020).

28. Eby, S. F. et al. Validation of shear wave elastography in skeletal muscle. J. Biomech. 46, 2381–2387 (2013).

29. Chernak, L. A., DeWall, R. J., Lee, K. S. & Thelen, D. G. Length and activation dependent variations in muscle shear wave speed. Physiol. Meas. 34, 713–721 (2013).

30. Wang, A. B., Perreault, E. J., Royston, T. J. & Lee, S. S. M. Changes in shear wave propagation within skeletal muscle during active and passive force generation. J. Biomech. 94, 115–122 (2019).

31. Gennisson, J.-L. et al. Viscoelastic and anisotropic mechanical properties of in vivo muscle tissue assessed by supersonic shear imaging. Ultrasound Med. Biol. 36, 789–801 (2010).

32. Sharafi, B. & Blemker, S. S. A micromechanical model of skeletal muscle to explore the effects of fiber and fascicle geometry. J. Biomech. 43, 3207–3213 (2010).

33. Witzenburg, C. M. & Barocas, V. H. A nonlinear anisotropic inverse method for computational dissection of inhomogeneous planar tissues. Comput. Methods Biomech. Biomed. Engin. 19, 1630–1646 (2016).

